# Miniaturized wireless bioelectronics for electrically driven biohybrid robots

**DOI:** 10.64898/2026.03.09.710657

**Authors:** Hiroyuki Tetsuka, Jiaju Ma, Minoru Hirano

## Abstract

Although biohybrid robots offer the potential for soft, adaptive actuation by harnessing living muscle, practical operation in cell culture environments is often limited by the requirement of immersed leads or cumbersome stimulation equipment. Here, we present a thin, miniaturized, wireless bioelectronic stimulator that can electrically drive biohybrid robots while maintaining stability in aqueous cell culture media. Built on a 50-µm liquid crystal polymer (LCP) substrate, the device integrates a planar receiving coil, interconnects, a diode-based rectifier, and a tank capacitor. This enables the device to convert an approximately 4.9-MHz radio-frequency (RF) input into pulsed direct current (DC), which is delivered through integrated stimulation electrodes. The stimulator has a footprint of ∼23 mm^2^ and a total thickness and mass of ∼100 µm and ∼7 mg, respectively. We integrated the stimulator with a nanopatterned carbon nanotube (CNT)/gelatin hydrogel fin seeded with human induced pluripotent stem cell-derived cardiomyocytes (iPSC-CMs) to generate propulsion through fin flapping. By optimizing the thickness of the polydimethylsiloxane (PDMS) encapsulation layer, the density was tuned, and the robot remained freely floating and retained shape integrity during operation. This produced autonomous forward locomotion of ∼70 µm/s. The stimulator generated distance-dependent output voltage pulses of ∼2–6 V and reliably synchronized fin flapping rates of up to 2 Hz without an observable loss of cell attachment or sarcomeric organization. Together, these results establish a compact, media-compatible, wireless, bioelectronic interface suitable for closed-system biohybrid robotics.

## Introduction

Biohybrid robots integrate living tissues with engineered structures to achieve lifelike actuation and biological autonomy that are difficult to replicate using conventional robotic systems ^1^. Such living muscle-based biohybrid platforms efficiently convert biochemical energy into motion and exhibit self-healing and adaptive responses, enabling life-like swimming, walking, and grasping behaviors ^2-21^. Despite these advancements, achieving reliable control of muscle contraction remains a central challenge for closed-system biohybrid robotics.

The induction of muscle contraction in skeletal and cardiac tissues, as well as regulation of contraction or beating rate, has relied primarily on electrical stimulation or optical stimulation in cell systems engineered to express light-responsive ion channels. However, both approaches typically require electrode wires or light sources such as optical fibers to be inserted within the culture medium. Wiring may hinder free motion of robots and complicates deployment in sealed vessels. Immersed components can perturb the local environment. For these reasons, a range of wireless bioelectronic approaches has been explored to provide stimulation without direct physical connections through the medium ^13,16,20,21^. In our earlier work, we built biohybrid platforms with wireless stimulators on polyurethane or polyimide substrates to pace cardiac tissues or excite motor nerve interfaces ^20,21^. These earlier platforms were comparatively large, with surface areas of at least 250 mm^2^. Also, they exhibited material-related limitations. Density mismatch complicated buoyancy and hydrodynamic tuning, while substrate deformation caused by moisture uptake impaired shape retention during long-term culture.

Here, we address these limitations by engineering a downsized wireless stimulation module (Fig. 1A). The stimulator is implemented on a liquid crystal polymer (LCP) substrate. We selected LCP for its low hygroscopicity and high dimensional stability. We build the inductive receiver and radio frequency (RF) to direct current (DC) conversion circuitry directly on LCP. Theor footprint is ∼23 mm^2^. This miniaturized platform is further integrated with a cardiomyocyte-powered carbon nanotube (CNT)/gelatin fin, where PDMS encapsulation is used to tune the apparent density toward that of the culture medium. Together, these designs yield a freely floating, electrically addressable biohybrid robot whose beating and fin motion can be paced wirelessly at frequencies of up to 2 Hz. This system provides a compact and robust route to closed-system biohybrid robotics.

**Figure 1.**
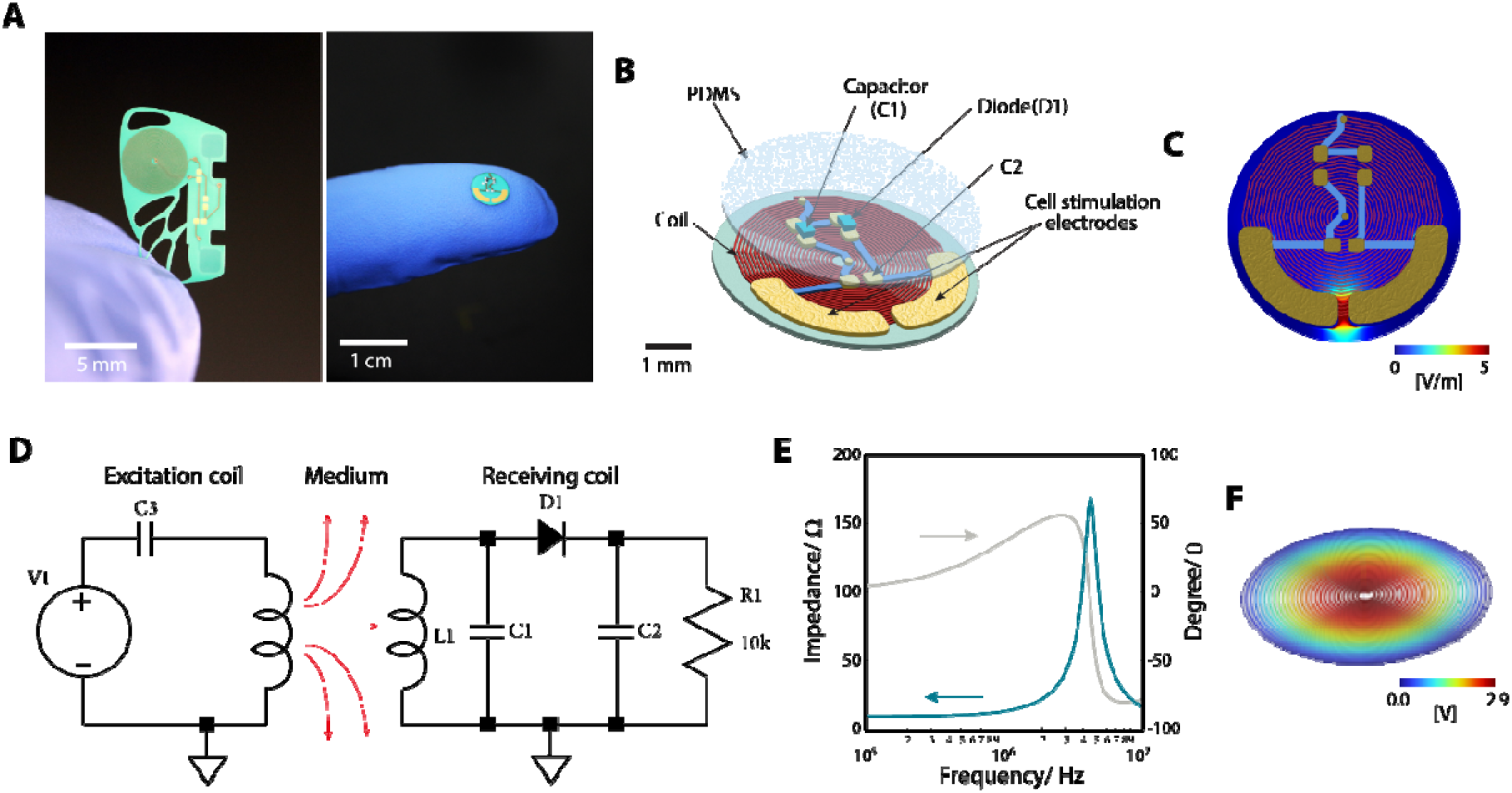
Miniaturized wireless bioelectronics. (A) Photograph of miniaturized wireless bioelectronics. (B) Layer-by-layer structure of the circular wireless device. (C) Electric field distribution in the device. (D) Schematic of the wireless system. (E) Resonant frequencies for the coil. (F) Electric potential distribution in the receiving coil.

## Results and Discussion

The miniaturized wireless bioelectronic stimulation module was realized on a 50 µm thick LCP substrate and integrates a planar receiving coil, stimulation electrodes, and rectification components, including a diode and a capacitor (Fig. 1A and 1B). The miniaturized planar receiving coil increases the design flexibility of the device, enabling a wide range of geometries, such as butterfly wing–inspired shapes and circular designs. In the equivalent circuit (Fig. 1D), the RF voltage induced in the receiver coil is converted into a pulsed DC waveform by a diode-based rectifier (D1) together with a capacitor (C1). The resulting output is delivered to the cell-facing electrodes, where the tissue load is represented as a resistor (R1). To ensure stable operation under RF excitation, the coil geometry (L1) was designed and evaluated using full-wave electromagnetic simulations. This approach enabled estimation of the inductive value near the operating frequency and verification that the self-resonant frequency (∼165 MHz; Fig. S1) was sufficiently separated from the excitation band (Fig. 1E). In addition, the receiving coil was confirmed to generate an induced voltage of 1 V or higher, which is required to stimulate muscle cells, as shown by the electric potential distribution at the transfer resonance frequency (Fig. 1F). Field mapping across the stimulation region further indicated that the potential gradient was concentrated around the interelectrode gap (Fig. 1C), suggesting that excitation would initiate near the electrodes and subsequently spread through the tissue via intercellular conduction.

We next assembled a biohybrid robot by integrating a CNT/gelatin hydrogel fin as the compliant actuator and seeding human iPSC-derived cardiomyocytes on its surface using the circular devices (Fig. 2A). LCP was selected as the structural base because of its low water uptake and high dimensional stability, which are advantageous for long-term operation under aqueous environment. The fabricated stimulator was ∼100 µm thick with a mass of ∼7 mg, and the actuator layer consisted of a CNT/gelatin hydrogel (∼150 µm thick) integrated onto the device (Fig. 2A). To protect the circuitry while simultaneously adjusting the overall density, the thickness of the PDMS encapsulation was tuned. The intrinsic density of LCP (∼1,300 kg m^−3^) lies between those of polyurethane (∼1,100 kg m^−3^) and polyimide (∼1,500 kg m^−3^) used previously, enabling more straightforward density matching using PDMS (density ∼900 kg m^−3^). With a PDMS coating of ∼400 µm, the apparent device density was estimated to be ∼937 kg m^−3^, close to that of the culture medium (∼1,000 kg m^−3^). Under these conditions, the assembled robot remained freely suspended in the medium without sinking or adhering to the vessel surfaces (Fig. S2).

**Figure 2.**
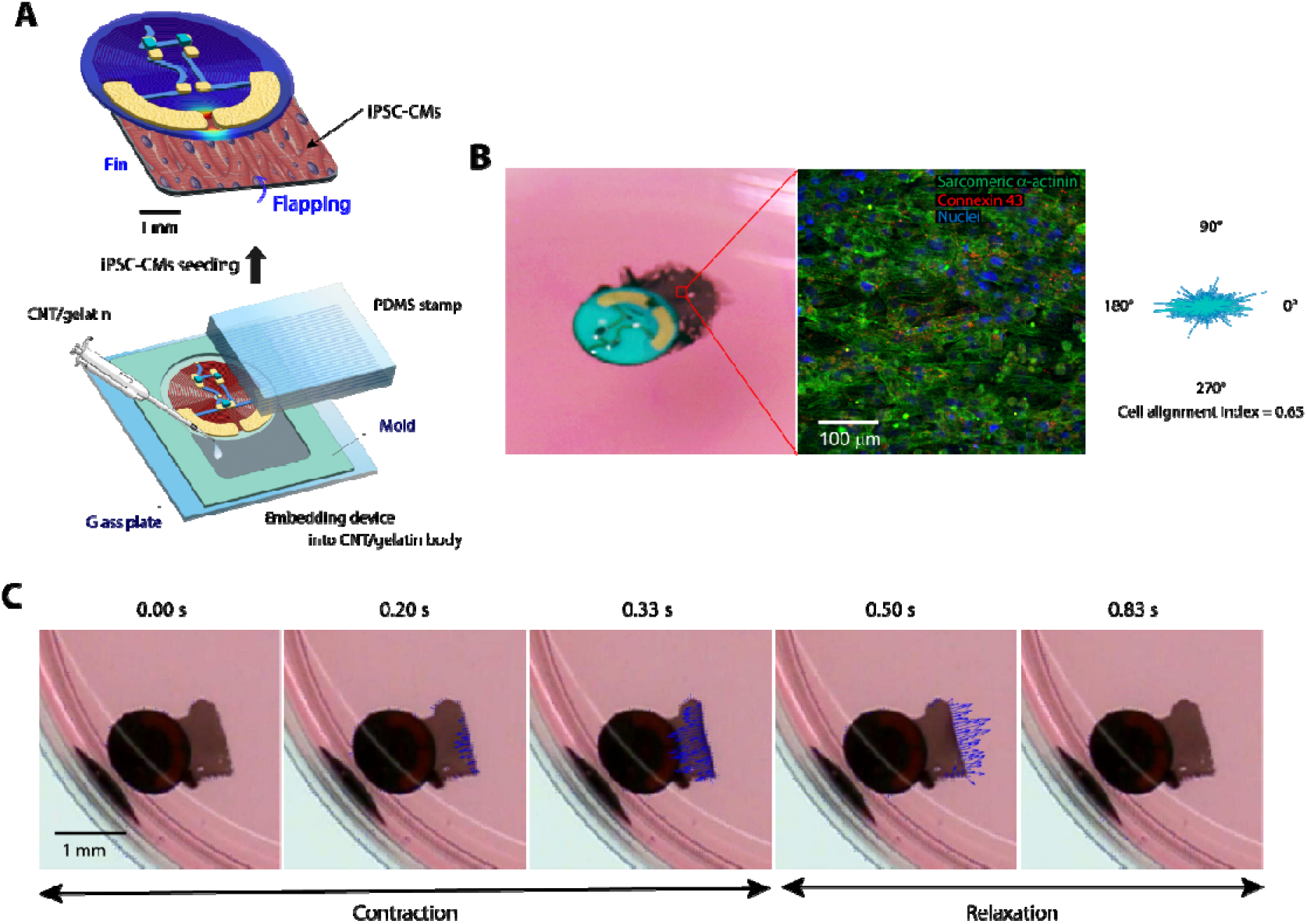
Integration of the wireless bioelectronics into biohybrid robots. (A) Fabrication process of bioelectronic robots. (B) Photograph of a bioelectronic robot released in cell culture medium and a confocal image of immunostained iPSC-derived cardiomyocytes on the fin. (C) Optical-flow analysis of the robot fin. The blue arrows show an alternating pattern of upward and downward motion vectors.

In this architecture, contraction timing is dictated by stimulation pulses delivered to the on-board electrodes through inductive power transfer, allowing the beating rate of the cardiomyocytes, and thus fin motion, to be set externally. A photograph of the complete robot and a microscope image of the fin region are shown in Fig. 2B; the latter confirms robust cardiomyocyte attachment to the CNT/gelatin scaffold. To promote anisotropic force generation, the CNT/gelatin surface was nanopatterned during enzymatic crosslinking using a grooved PDMS stamp with a groove width and depth of 0.8 µm. This patterning biased cellular alignment along the groove direction, producing a tissue that contracts preferentially along a single axis. During contraction, the aligned tissue shortens and drives the hydrogel to bend upward, while relaxation restores the fin toward its undeformed configuration. Repeated cycles of contraction and relaxation therefore generate vertical flapping, which translates into forward thrust.

Kinematic quantification based on video analysis indicated that, during spontaneous beating, the fin reached peak upward deformation in ∼0.33 s and then relaxed over ∼0.50 s (Fig. 2C). The resulting free-swimming motion is summarized in Fig. S3: the robot advanced predominantly forward with an average speed of ∼70 µm s^−1^. This performance was comparable to that of previously reported systems integrating a wireless stimulator on a polyurethane substrate, which achieved an average speed of ∼24 µm s−1 (Fig. S3) ^20^.

Wireless pacing characteristics were evaluated by transmitting the stimulation waveforms shown in Fig. 3A and recording the output from the device. For example, with a pulse width of 50 ms and a repetition rate of 1 Hz, the module produced an output of ∼2 V when positioned 1 cm from the transmitting coil (Fig. 3A and 3B). The output amplitude increased as the separation distance decreased, reaching ∼6 V when the device was placed close to the transmitter. The fin response to externally applied pacing is summarized in Fig. 4. In the absence of stimulation, the fin beat spontaneously at ∼0.7 Hz. When a 1 Hz pulse train was applied, the flapping rate shifted to 1.0 Hz and remained synchronized with the imposed rhythm (Fig. 4B). After stimulation was halted, the fin returned to its spontaneous activity at ∼0.7 Hz. The system also followed higher-frequency commands, with stable pacing observed for input frequencies of up to 2 Hz (Fig. 4C). Together, these observations demonstrate that the miniaturized module can impose user-defined contraction timing on the cardiomyocyte actuator and thereby modulate swimming dynamics through a purely wireless link.

**Figure 3.**
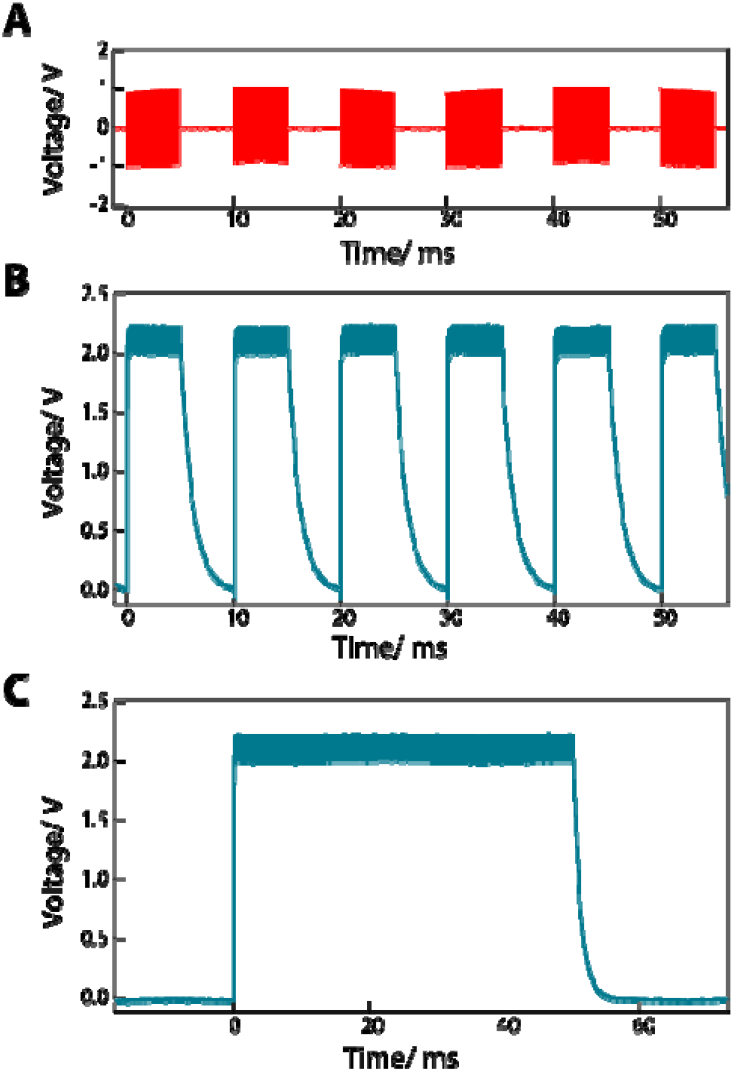
Wireless electrical stimulation signals. (A) Example of a waveform. Rectangular pulses with a width of 50 ms and a frequency of 1 Hz were repeated at an interval of ∼1 s. (B) Generated signals from the wireless device using the waveform shown in (A). (C) Expanded plot of (B).

**Figure 4.**
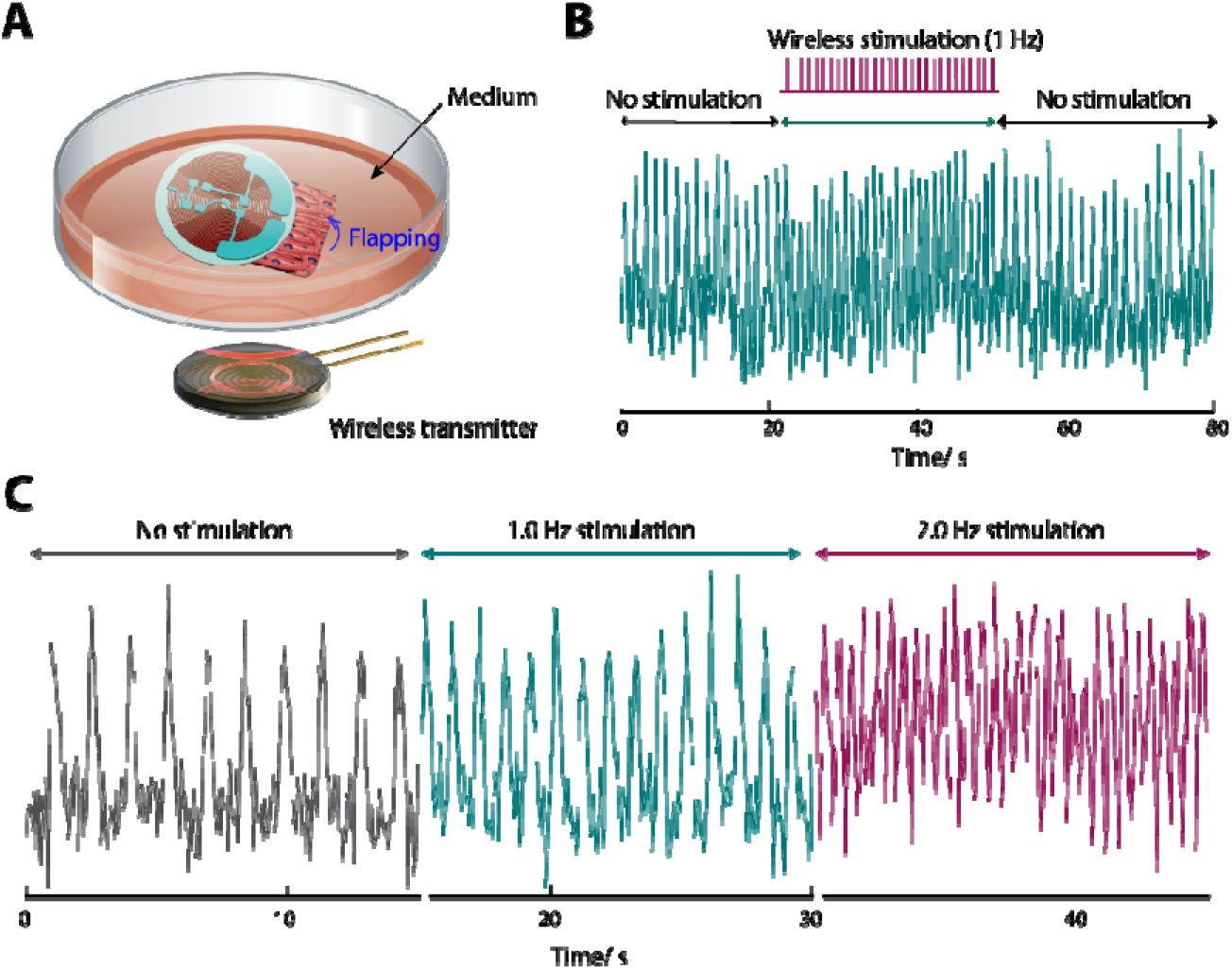
Wireless control of bioelectronic robots. (A) Setup for wireless stimulation of robots. (B) Representative temporal trace of fin deflection during the wireless 1 Hz electrical stimulation. (C) Temporal trace of fin deflection under different wireless activates frequencies. All deflection behaviors and controllability were analyzed for robots that were placed and secured on a dish in the medium to eliminate the influence of forward locomotion.

Finally, tissue integrity after stimulation was assessed by immunostaining (Fig. S4). No obvious detachment of cells from the fin surface was observed after wireless operation (Fig. S4A), and a cardiac tissue layer with a thickness of ∼30 µm remained attached (Fig. S4B). On the nanopatterned CNT/gelatin fin, cardiomyocytes exhibited pronounced single-direction alignment and retained sarcomeric features characteristic of striated muscle, including α-actinin striations (Fig. S4C).

## Conclusion

We engineered a thin, lightweight wireless bioelectronic interface that enables biohybrid robots to be paced and actuated without immersed wiring. The device, fabricated on a 50 µm LCP substrate, integrates an inductive receiver and rectification elements, including a diode and a capacitor, to convert a ∼4.9 MHz RF signal into stimulation DC pulses delivered through on-board electrodes. Miniaturization reduced the active footprint to ∼23 mm^2^ while maintaining a total thickness of ∼100 µm and a mass of ∼7 mg, thereby mitigating the mechanical and buoyancy limitations associated with larger prior platforms. Coupling the stimulator to a nanopatterned CNT/gelatin hydrogel fin seeded with human iPSC-derived cardiomyocytes produced repeatable fin deformation and forward swimming. Encapsulation with a tuned PDMS layer adjusted the apparent density toward that of the culture medium, supporting stable free-floating operation and preserving shape retention in aqueous environments. The wireless stimulator generated ∼2–6 V pulses depending on transmitter distance and provided reliable pacing control, shifting spontaneous fin activity (∼0.7 Hz) to externally defined rates of up to 2 Hz. Post-stimulation assessments indicated maintained tissue attachment and preserved muscle organization. Together, these results establish a compact, media-compatible wireless bioelectronics approach for electrically driven biohybrid machines, opening a path toward more versatile closed-system biohybrid robotics and scaling to smaller, multi-actuator, or more complex systems.

## Methods

### Fabrication of wireless bioelectronic cellular stimulation devices

Miniaturized wireless bioelectronic cellular stimulation devices were manufactured by an external vendor (Yamashita Material Co., Ltd.). After delivery, the devices were encapsulated in PDMS to provide electrical insulation and environmental protection.

### Fabrication of biohybrid robots

#### Materials

Gelatin from porcine skin (Type A, ∼300 g bloom) was obtained from Sigma–Aldrich. A suspension of carboxylic acid–functionalized multiwalled CNTs was provided by Hamamatsu Carbonics Inc. Microbial transglutaminase (Moo Gloo TI) was obtained from Modernist Pantry Inc. Dulbecco’s phosphate-buffered saline (DPBS) was sourced from Thermo Fisher Scientific Inc. Human iPSC-derived cardiomyocytes (single-cell suspension), along with the associated plating and maintenance media, were obtained from FUJIFILM Cellular Dynamics, Inc. (iCell Cardiomyocytes^2^ Kit).

#### Integration of CNT/gelatin fin into wireless devices

A wireless stimulator was positioned on a glass support and aligned using a molding fixture. A CNT/gelatin precursor solution (0.1 w/v% CNT and 10 w/v% gelatin) was combined with microbial transglutaminase (mTG; 10 U per gram of gelatin), and 100 µL of the mixture was dispensed into the mold covering the device. To introduce surface nanotopography and define film thickness, a PDMS stamp with a groove width and depth of 0.8 µm was placed on the hydrogel surface immediately after casting. A 100 g calibration weight was then applied to compress the gel and remove excess precursor, yielding the targeted thickness. The construct was left to crosslink enzymatically overnight at room temperature under a glass cover to minimize drying. The weight was removed after gelation.

#### Sterilization protocol

Following scaffold formation, constructs were rinsed with DPBS supplemented with 10% antibiotic–antimycotic solution (Thermo Fisher Scientific Inc.) and incubated for 10 min at room temperature. The constructs were then transferred to DPBS containing 1% antibiotic–antimycotic solution and maintained overnight at 37°C prior to cell seeding.

#### Culture of iPSC-CMs

After completion of the sterilization procedure, iPSC-derived cardiomyocytes were seeded onto each construct as a single-cell suspension at a density of ∼2.5 × 105 cells/cm^2^ in plating medium. Cells were allowed to attach for 4 h at 37°C, after which the medium was replaced with maintenance medium. Maintenance medium was refreshed every other day thereafter.

### Immunostaining for cell characterization

Biohybrid robots were washed with DPBS and fixed in 4% paraformaldehyde for 15 min. Fixed samples were rinsed three times with DPBS and then incubated for 15 min at room temperature in a blocking solution consisting of DPBS containing 0.2% Triton X-100 and 10% goat serum. Primary antibodies (mouse anti-sarcomeric α-actinin and rabbit anti-connexin 43) were diluted 1:200 in blocking buffer and applied to samples overnight at 4°C. After three DPBS washes, samples were incubated for 30 min with secondary antibodies (goat anti-mouse IgG-AF488 and goat anti-rabbit IgG-AF594), each diluted 1:200 in blocking buffer. Nuclei were counterstained with DAPI (1:1,000 in DPBS) for 15 min. Fluorescence images were acquired using an inverted fluorescence microscope and a laser-scanning microscope.

### Cell alignment index

Alignment of sarcomeric structures was quantified from α-actinin immunofluorescence images using a custom MATLAB (MathWorks, Inc.) script. Briefly, two-dimensional Fourier power spectra were computed from each image, transformed into polar coordinates with 1° angular resolution, and averaged as a function of angle. Eigenvalues of the resulting orientation matrix were used to calculate an orientation, or alignment, index, where 0 denotes random organization and 1 denotes perfect uniaxial alignment.

### Evaluation of autonomous propulsion of biohybrid robots

Robot motion was recorded under a microscope equipped with a digital camera. Trajectories were extracted from videos using Tracker (http://physlets.org/tracker) by determining robot positions frame by frame. In addition, a custom MATLAB routine using the Computer Vision Toolbox was employed for motion vector analysis. Multiple digital markers were assigned to each robot based on pixel intensity features. Displacements between consecutive frames were computed to obtain local velocities and directions, defined as the magnitude and angle of the frame-to-frame displacement vectors, respectively.

### Optical flow analysis

Optical flow analysis was implemented using a custom MATLAB pipeline with the Computer Vision Toolbox. Feature points were identified on the robots based on image intensity patterns and tracked across successive frames. For each feature point, the displacement between frames was calculated, with speed defined as the displacement magnitude and directionality defined by the displacement angle.

### Electromagnetic simulation of wireless bioelectronic cellular stimulation devices

Electric field distribution within the wireless device was computed using the finite element method (FEM) in MATLAB with the Partial Differential Equation Toolbox. A three-dimensional geometry of the device was imported directly from a stereolithography (STL) file and used to define the computational domain. Electric potential distribution in the receiving coil was simulated using FEM in COMSOL Multiphysics (COMSOL, Inc.).

## Author contributions statement

H.T. conceived and designed the project. H.T. and J.M. designed the robot. H.T. conducted experiments, analyzed data, and wrote the manuscript. J.M. designed, fabricated, and characterized the wireless system. J.M. and M.H. edited the manuscript. All authors discussed the results and commented on the manuscript.

## Additional information

The authors declare no conflict of interest. All data required to evaluate the conclusions in the paper are presented in the main manuscript and or the Supplementary Materials. Additional data are available from the authors upon request.

