## Supplemental figures S1-4 for "Miniaturized wireless bioelectronics for electrically driven biohybrid robots"

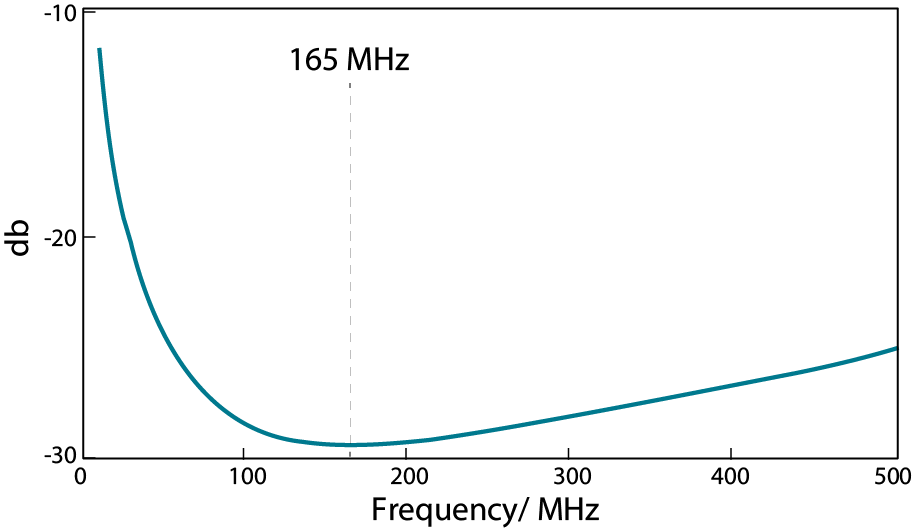

Fig. S1. Self-resonant frequency of the receiving antenna coil.

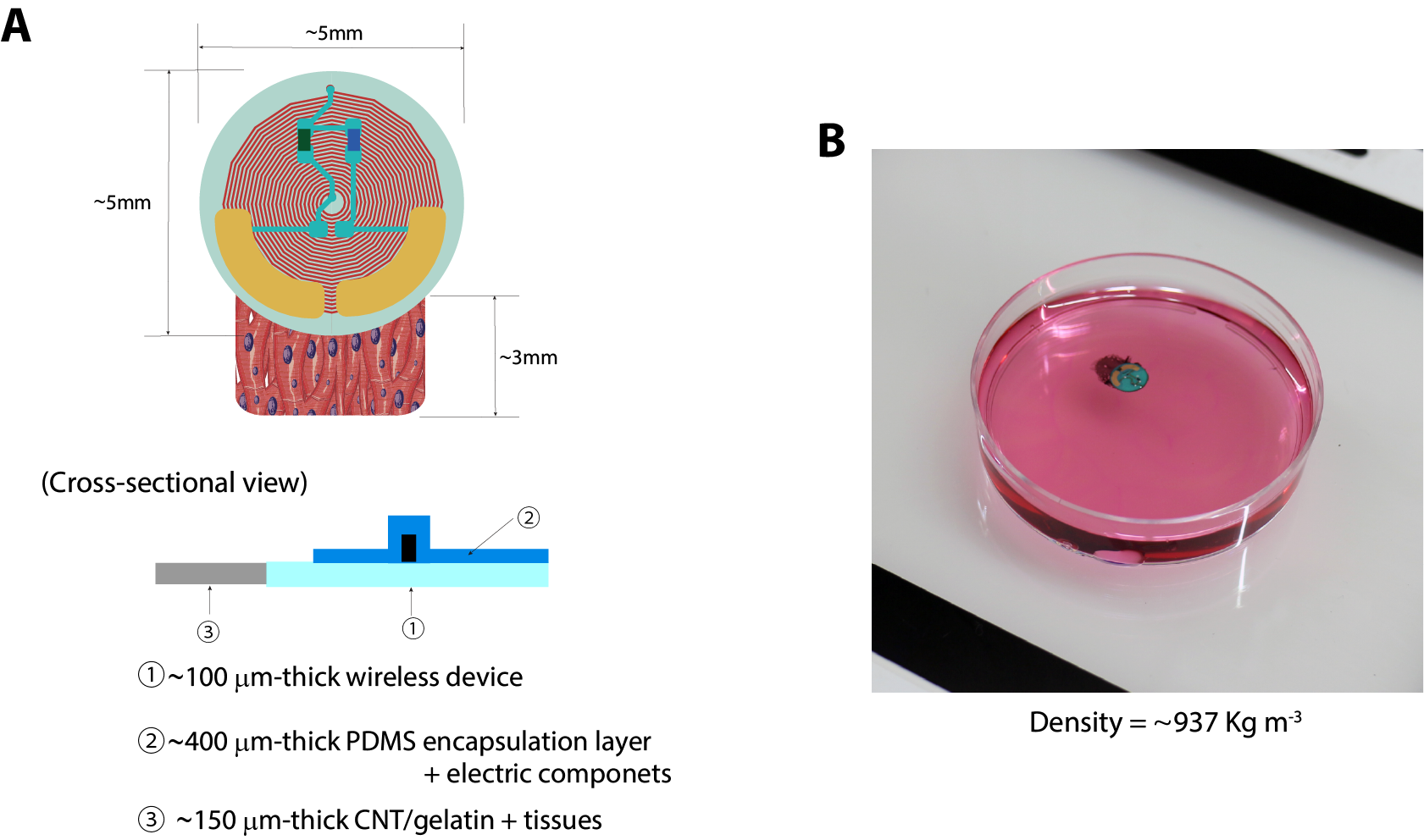

Fig. S2. (A) Design and cross-sectional layer-by-layer structure of a bioelectronic robot. (B) Photograph of a bioelectronic robot released in the cell culture medium.

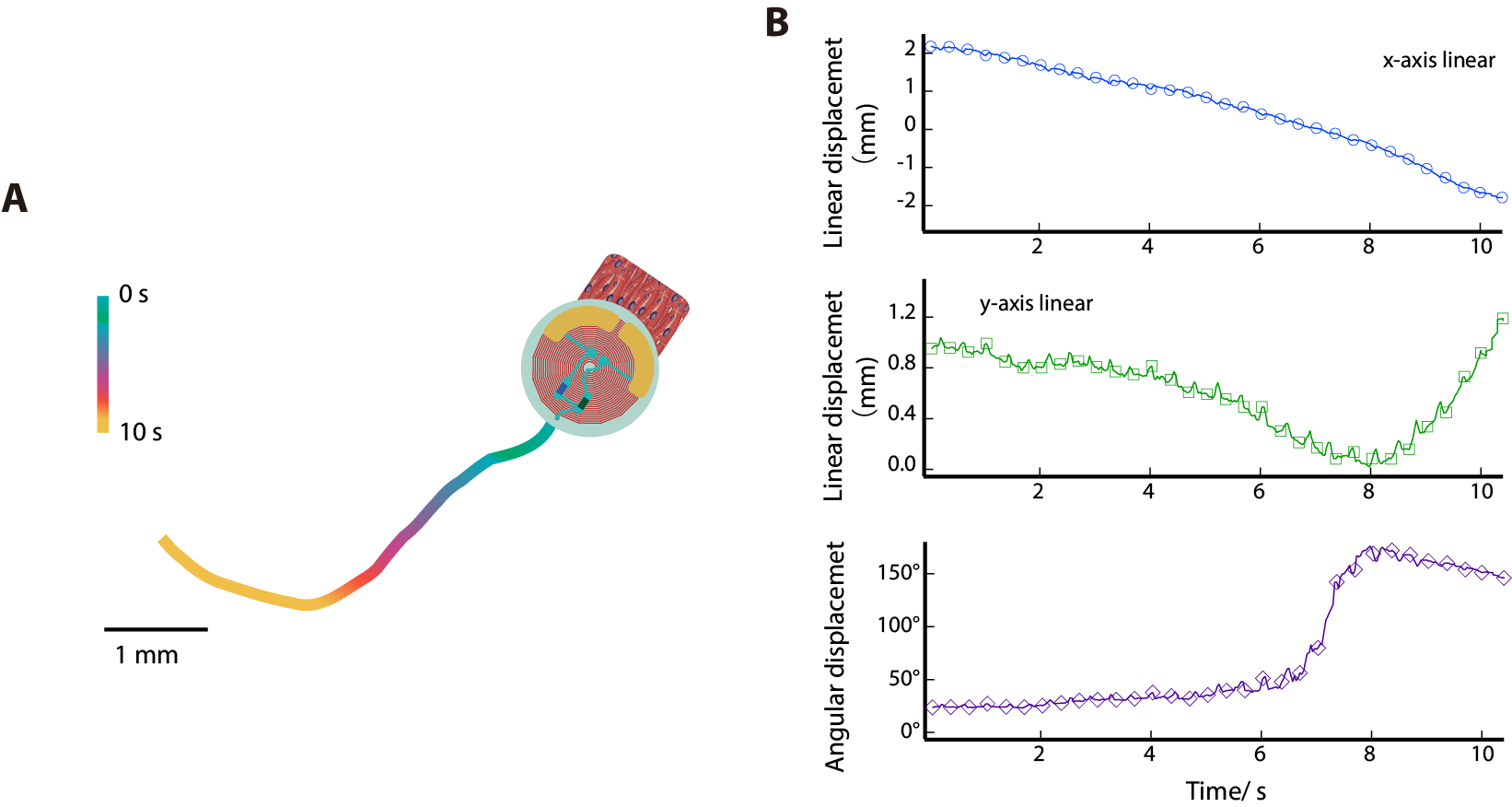

Fig. S3. (A) Locomotion trajectory of a bioelectronic robot. (B) Linear and angular displacements of (A).

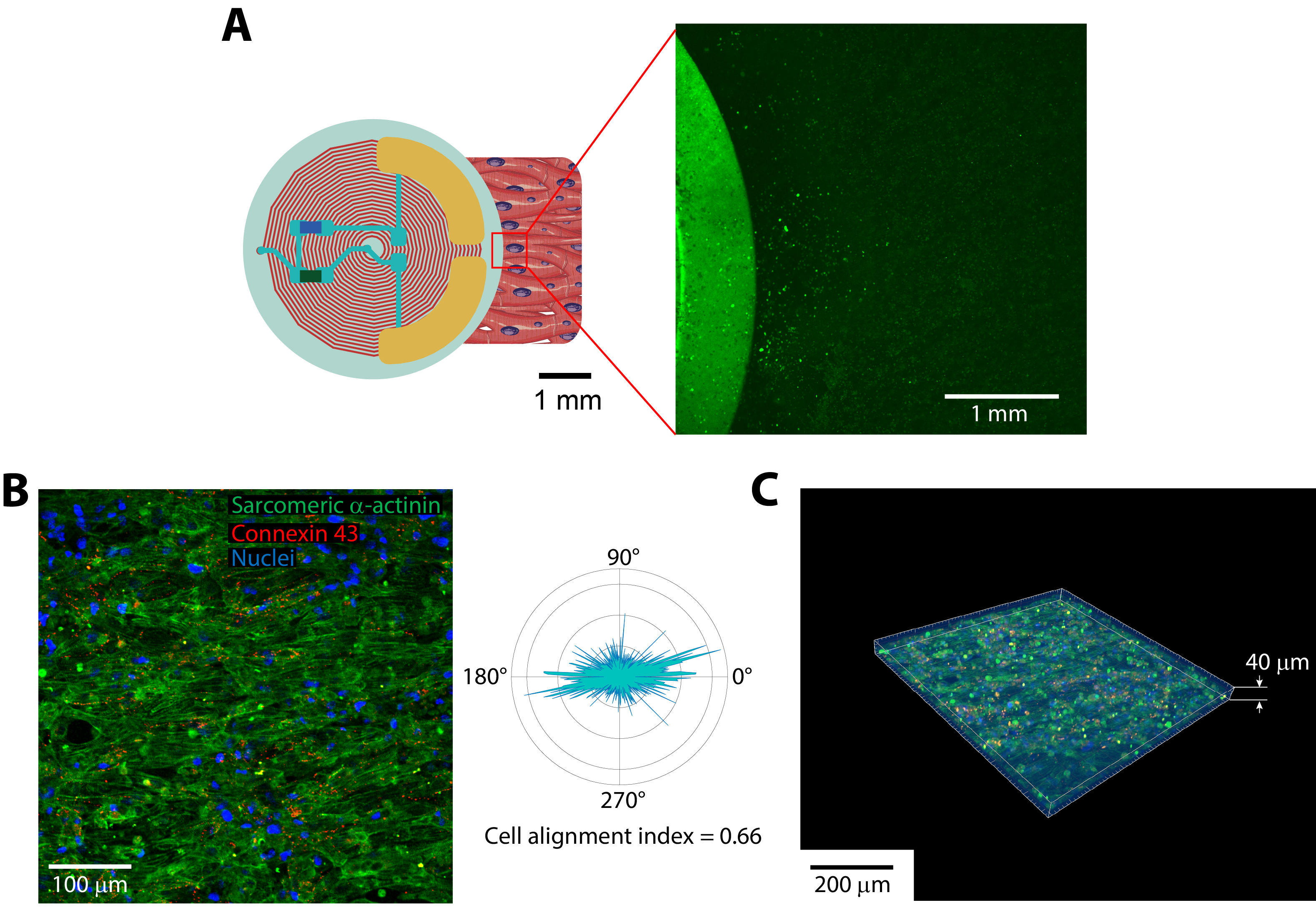

Fig. S4. Fluorescence (A) and confocal (B, C) microscopic images of the robot after wireless cell stimulation trains.
